# Single cell-guided prenatal derivation of primary epithelial organoids from the human amniotic and tracheal fluids

**DOI:** 10.1101/2023.05.31.539801

**Authors:** Mattia Francesco Maria Gerli, Giuseppe Calà, Max Arran Beesley, Beatrice Sina, Lucinda Tullie, Francesco Panariello, Federica Michielin, Kylin Sun Yunyan, Joseph R Davidson, Francesca Maria Russo, Brendan C Jones, Dani Lee, Savvas Savvidis, Theodoros Xenakis, Ian Simcock, Anna A Straatman-Iwanowska, Robert A Hirst, Anna L David, Christopher O’Callaghan, Alessandro Olivo, Simon Eaton, Stavros P Loukogeorgakis, Davide Cacchiarelli, Jan Deprest, Vivian SW Li, Giovanni Giuseppe Giobbe, Paolo De Coppi

**Affiliations:** UCL Department of Surgical Biotechnology, Division of Surgery and Interventional Science, London, UK; UCL Great Ormond Street Institute of Child Health, London, UK; Politecnico di Milano, Milano, Italy; Stem Cell and Cancer Biology Laboratory, The Francis Crick Institute, London, UK; Telethon Institute of Genetics and Medicine, Armenise/Harvard Laboratory of Integrative Genomics, Pozzuoli, Italy; UCL Elizabeth Garrett Anderson Institute for Women’s Health, London, London, UK; UZ Leuven Clinical Department of Obstetrics and Gynaecology, Leuven, Belgium; UCL Department of Medical Physics and Biomedical Engineering, London, UK; Department of Radiology, Great Ormond Street Hospital, London, UK; University of Leicester, Department of Respiratory Sciences, Leicester, UK; Specialist Neonatal and Paediatric Surgery, Great Ormond Street Hospital for Children NHS Foundation Trust, London, UK

## Abstract

Despite advances in prenatal diagnosis, it is still difficult to predict severity and outcomes of many congenital malformations. New patient-specific prenatal disease modelling may optimise personalised prediction. We and others have described the presence of mesenchymal stem cells in amniotic fluid (AFSC) that can generate induced pluripotent stem cells (iPSCs). The lengthy reprogramming processes, however, limits the ability to define individual phenotypes or plan prenatal treatment. Therefore, it would be advantageous if fetal stem cells could be obtained during pregnancy and expanded without reprogramming. Using single cell analysis, we characterised the cellular identities in amniotic fluid (AF) and identified viable epithelial stem/progenitor cells of fetal intestinal, renal and pulmonary origin. With relevance for prenatal disease modelling, these cells could be cultured to form clonal epithelial organoids manifesting small intestine, kidney and lung identity. To confirm this, we derived lung organoids from AF and tracheal fluid (TF) cells of Congenital Diaphragmatic Hernia (CDH) fetuses and found that they show differences to non-CDH controls and can recapitulate some pathological features of the disease. Amniotic Fluid Organoids (AFO) allow investigation of fetal epithelial tissues at clinically relevant developmental stages and may enable the development of therapeutic tools tailored to the fetus, as well as to predicting the effects of such therapies.

## INTRODUCTION

Modern prenatal screening routinely adopts sophisticated genetic and imaging analyses that are increasingly effective at detecting and characterising congenital anomalies^1, 2^. Despite this, it remains challenging to predict the functional severity of many complex conditions such as CDH, spina bifida (SB), cystic fibrosis (CF) and polycystic kidney disease (PKD) after their identification, patient-specific parental counselling is limited. Stratification of prenatal therapy is very relevant and has been used in the last few years to select patients for fetal intervention that may reverse the natural history of some of these conditions. There is level 1 evidence for improved outcomes in conditions such as CDH^3, 4^ and SB^5^, while for other conditions such as in vesico-amniotic shunting for lower urinary tract obstruction (LUTO)^6^ is technically possible, but appropriate patient selection remains the main hurdle. The lack of autologous systems that are capable of accurately recapitulating the complexity and functional characteristics of developing human tissues is the main bottleneck to implementing personalised regenerative medicine strategies that would deliver a significant impact in the lives of babies affected by these conditions. Consequently, in terms of directing prenatal therapy, the field of prenatal functional diagnosis remains underdeveloped.

Organoids provide a reliable three-dimensional tissue model, which can recapitulate some of the biological and pathophysiological features of the patient’s tissues *in vitro*. Autologous organoids can be derived from human embryonic stem cells^7^ or during prenatal and postnatal life through reprogramming to iPSCs^8^. These cells are then committed to the specific tissue-type of interest and expanded into stable lines^9–14^. With respect to human developmental conditions, iPSC-derived organoids have been successfully generated from fetal cells within the AF of both healthy fetuses and those with congenital anomalies^15, 16^. Overall, human iPSC-derived organoids have some advantages in terms of patient-specificity, but their reliance on considerable manipulation reduces fidelity to the patient’s individual condition, and strict quality control bears significant cost and time implications that hamper their applicability to personalised disease modelling in order to target therapy.

Primary organoids have previously been derived from numerous human tissues^17–20^. Being directly derived from the target tissues, these require significantly fewer *in vitro* manipulations, hence carrying lower safety burdens. More recently, primary organoids have been derived from discarded postnatal biological samples (e.g., urine, menstrual flow, PAP brush, bronchoalveolar lavage^20–23)^. This has clear advantages over the use of biopsies, allowing generation of autologous primary organoids from samples that are anyway acquired in the course of routine clinical care. In the context of prenatal medicine, primary organoids have been successfully derived from several fetal tissues collected post-mortem or through biobanks such as the Human Developmental Biology Resource (HDBR)^24–26^. The use of primary fetal organoids offers many unprecedented benefits to study human development and congenital diseases. Access to fetal tissues, however, has ethical and legal restrictions in many parts of the world that prohibit their use in research^27^. Until now, primary fetal organoids have only been derived with destructive methods, limiting their use for autologous disease modelling, prenatal functional diagnostics and personalised therapeutics^28^. In this manuscript, we present the derivation of primary human fetal epithelial organoids of multiple tissue identities (intestinal, renal and pulmonary) from fetal fluids collected during the second and third trimester of gestation. The fluids are already sampled as part of routine prenatal diagnosis and therapeutic intervention. The process permits the generation of organoids alongside the continuation of the pregnancy.

The AF surrounds, supports and protects the human fetus during development. The origin and recirculation of this fluid follows complex dynamics, evolving together with the development of the various fetal and extraembryonic tissues that contribute to its production. As consequence, the AF contains cells shed from a variety of origins^29^. The AF harbours multipotent stem cells first identified by ours and other teams and since ascribed to the mesenchymal and haematopoietic niches^30–32^. AFSCs have successfully been used in various animal models of disease, where they have demonstrated the capacity to induce regeneration through cell transplantation, conditioned media and activated AFSC-derived extracellular vescicles^33–36^. Whilst stem cells with mesenchymal and hematopoietic potential have clear therapeutic promise, most of the cells present in the AF manifest an epithelial identity. The AF is highly heterogeneous in origin and composition and includes secretions and cells from various tissues, including the fetal kidney, lung and gastrointestinal tract^29, 37^. Due to the complexity of the epithelial culture systems, a detailed map of the AF epithelial population has not yet been compiled. Thanks to advances in the fields of single cell sequencing and organoid biology, it is now feasible to investigate these possibilities in developing tissues^38^. With these techniques, we validated the presence of amniotic fluid epithelial cells (AFEC), highlighting that these shed from a multiplicity of developing tissues, and demonstrating that this population contains progenitors capable of forming tissue-specific primary fetal organoids. Finally, with the adoption of fetal surgery procedures such as Fetal Endoluminal Tracheal Occlusion (FETO) to treat CDH, we can now sample and expand viable epithelial cells from tracheal fluid (TF) at various stages in the therapy ^39–41^.

In this work we provide evidence that diverse epithelial stem cell populations are shed from a number of different tissues into the AF and TF during development. We then show that these cells are capable of forming tissue-specific primary fetal organoids. Autologous derivation of primary fetal organoids during a continuing pregnancy would broaden the possibility of conducting research at later gestational stages which may be beyond the limits of termination of pregnancy, allowing the development of functional diagnosis, personalised counselling, and providing innovative tools for designing autologous prenatal and perinatal therapies.

## RESULTS

### Single cell mapping of the human amniotic fluid reveals the presence of fetal intestinal, renal and pulmonary epithelial progenitors

With the aim of mapping the cellular content of the human AF and investigating the presence of epithelial progenitors from multiple tissues, we collected AF from 11 pregnancies ranging between 15-34 weeks’ gestational age (GA) (Supplementary Table 1). As more than 97% of the cells shed in the AF are dead or undergoing apoptosis, we first isolated the viable cell fraction using fluorescence-activated cell sorting (FACS). To preserve the heterogeneity of the AF cellular content, we did not select the cells by Forward Scatter or Side Scatter. The incorporation of Hoechst was used to identify nucleated cells and exclude debris, urea crystals and other non-cellular particles. In conjunction, we used propidium iodide (PI) to exclude dead cells and cells with a damaged plasma membrane (Figure 1a; Supplementary Figure 1a). After confirming cell viability through Live/Dead Acridine Orange/Propidium Iodide fluorescent Luna cell counter, we performed 3’ single cell RNA sequencing (scRNAseq) using the 10x Genomics Chromium platform. The data were processed, batch corrected and filtered to produce normalised count matrices, enabling the generation of an unsupervised uniform manifold approximation and projection (UMAP) using Seurat v4 (Figure 1a middle and Supplementary Figure 1b). We applied the SingleR package to perform automated labelling of any AFEC based on human primary cell atlas data^42^. We then probed our dataset for the expression of the epithelial genes *EPCAM*, *CDH1*, *KRT8*, *KRT10, KRT17* and *KRT19*, which showed high levels of expression (Figure 1a right; Supplementary Figure 1c). We validated that the cluster expressing these epithelial markers was the same as highlighted by SingleR (Figure 1b). A further validation of these gene expression data was performed by flow cytometry on viable, unselected AF cells, confirming the broad presence of EpCAM (EPCAM) and ECAD (CDH1) in the majority of live cells present in the AF (Figure 1c). We proceeded to interrogate our scRNAseq data for the expression of distinct intestinal, renal, and pulmonary epithelial markers within the AFEC cluster. The violin plots presented in Figure 1d highlight the presence of specific markers of the three tissues within the AF epithelial cluster. Finally, we assessed expression of tissue-specific stem / progenitor cell markers in the dataset. The UMAPs highlight the presence of cells co-expressing: *SOX2 and ASCL2, MUC2, FABP1, LRIG1* (Intestine), *PAX8* or *LHX1* (kidney), and at least two of *SOX2,* SOX9, *NKX2*-1, *FOX2A* (Lung) (Figure 1e). This demonstrated the presence of intestinal, renal and pulmonary stem / progenitor cells in the AF. Based on this evidence, we went on to culture primary fetal organoids from the AF.

**Figure 1.**
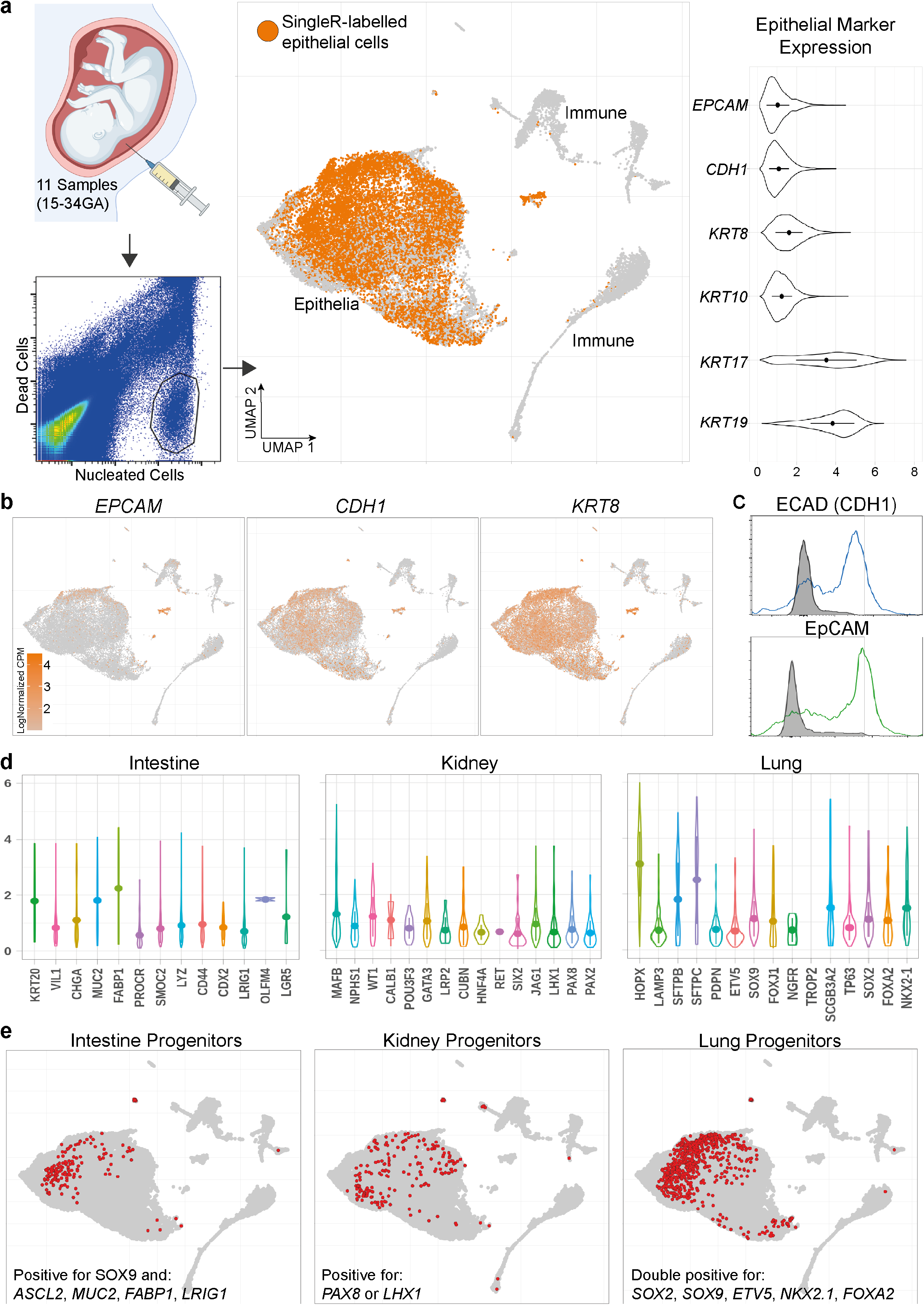
– Single cell analysis of the AF content. (**a**) Graphical representation of the AF sampling (top). The bottom plot shows the sorting strategy utilised to collect the living cell fraction, negative for Propidium Iodide and positive for Hoechst. The UMAP shows the content of the amniotic fluid of multiple patients obtained across the second trimester of pregnancy (n=11 patients 15-34GA; 40919 single cells are presented). The UMAP shows the content of the amniotic fluid of multiple patients obtained across the second trimester of pregnancy (n=11 patients 15-34GA; 36726 single cells analysed). Highlighted in orange the epithelial cluster, as identified by the SingleR cell labelling package using the human primary cell atlas dataset as reference. The violin plots show the level of expression of the pan-epithelial specific genes *EPCAM, CDH1(ECAD), KRT8, KRT10, KRT17* and *KRT19* (data presented as normalised counts per million, CPM). **(b)** The UMAPs show the expression of the same epithelial markers, within the epithelial cluster identified in **a**. **(c)** Representative flow cytometry analysis of EPCAM and ECAD (CDH1) expression in live-sorted cells from the AF, grey line represents unstained controls. **(d)** The violin plots highlight the occurrence, and level of expression of intestinal, renal and pulmonary cell markers within the epithelial cell cluster highlighted above (data presented as normalised CPM). (**e**) The UMAPs highlight in red, the presence of cells co–expressing tissue specific epithelial stem/progenitor cells markers: *SOX2 and ASCL2, MUC2, FABP2, LRIG1* (Intestine), *PAX8* or *LHX1* (kidney), and two of *SOX2, SOX9, NKX2*-1, *FOX2A* (Lung).

### Generation of primary fetal epithelial human amniotic fluid organoids (AFO)

We isolated unselected viable AF cells with the method described above. The cells were seeded in Matrigel droplets and cultured in an *ad hoc-*defined epithelial medium, developed to allow the formation of organoids, without providing specialised tissue-specific signals. Using this method, we derived >200 organoid lines from 18 AF samples ranging from 16-34 weeks GA (Supplementary Table 1). Organoid formation was observed in 82% of the AF samples (18/22), with a median formation efficiency of 9 organoids formed/10^5^ live cells plated (range 30.9). However, we found no meaningful association between the GA at AF sampling and the AFO formation efficiency (Figure 2b; supplementary figure 2a). Starting from day zero, some cells began proliferating and self-organising to form 3D structures, that became visible within the first week. During the first 2 weeks of culture, the formation of organoids became evident, and it was possible to pick single clonal organoids (Figure 2a-c). Once dissociated into single cells and replated, the organoids started reforming and expanding. AFO showed multiple morphological features, could be expanded at least until passage 5 and successfully cryopreserved, providing initial evidence of self-renewal and long-term culture (Figure 2b-d; Supplementary Figure 2b-c). Two distinct organoid phenotypes were observed through X-ray phase contrast computed tomography (PC-CT) and were determined to be parenchymatous and cystic (Figure 2e). To further confirm the presence of proliferating cells within the AFO, as well as the lack of apoptotic cell death, we performed immunostaining for the proliferative marker Ki67 and the apoptotic marker caspase 3 (Figure 2f). To confirm the epithelial identity of the organoids, we performed immunofluorescent staining for standard pan-epithelial human cell markers (EpCAM, ECAD, and Pan-Cytokeratin). Concomitantly, we have confirmed the lack of mesenchymal features by demonstrating absence of PDGF receptor alpha expression. Interestingly, AFOs form a polarised epithelium as demonstrated by the presence of basolateral integrin b4 (ITGβ4), apical F-Actin and the tight junction marker ZO-1 on the luminal side of the organoids (Figure 2g). To gather initial evidence on the tissue identity of the organoids, we isolated RNA from clonally expanded organoid lines and performed bulk RNA sequencing (n=105 from 17 independent AF samples). The bulk RNA sequencing data was subjected to an unsupervised principal component analysis (PCA; Figure 2h; Supplementary Figure 2d). Primary fetal organoids were derived from fresh fetal human intestine, stomach, lung and kidney, and included in the dataset as internal controls (n=14 with a minimum of 3 per tissue type). The PCA highlights the formation of three clusters of samples, colocalising with the intestinal, pulmonary and renal controls respectively (confirmed by Euclidean clustering, Supplementary Figure 2e). Moreover, a Gene Ontology analysis performed on each cluster against the rest of the dataset (after removal of the control organoids) showed upregulation of pathways related to the three tissues in the respective cluster (Figure 2i). Overall, this provides evidence that AF cells can give rise to clonal small intestine, lung and kidney AFO. We then went on investigating the gene and protein expression of tissue specific markers, as well as the organoids’ maturation potential within the three AFO clusters.

**Figure 2.**
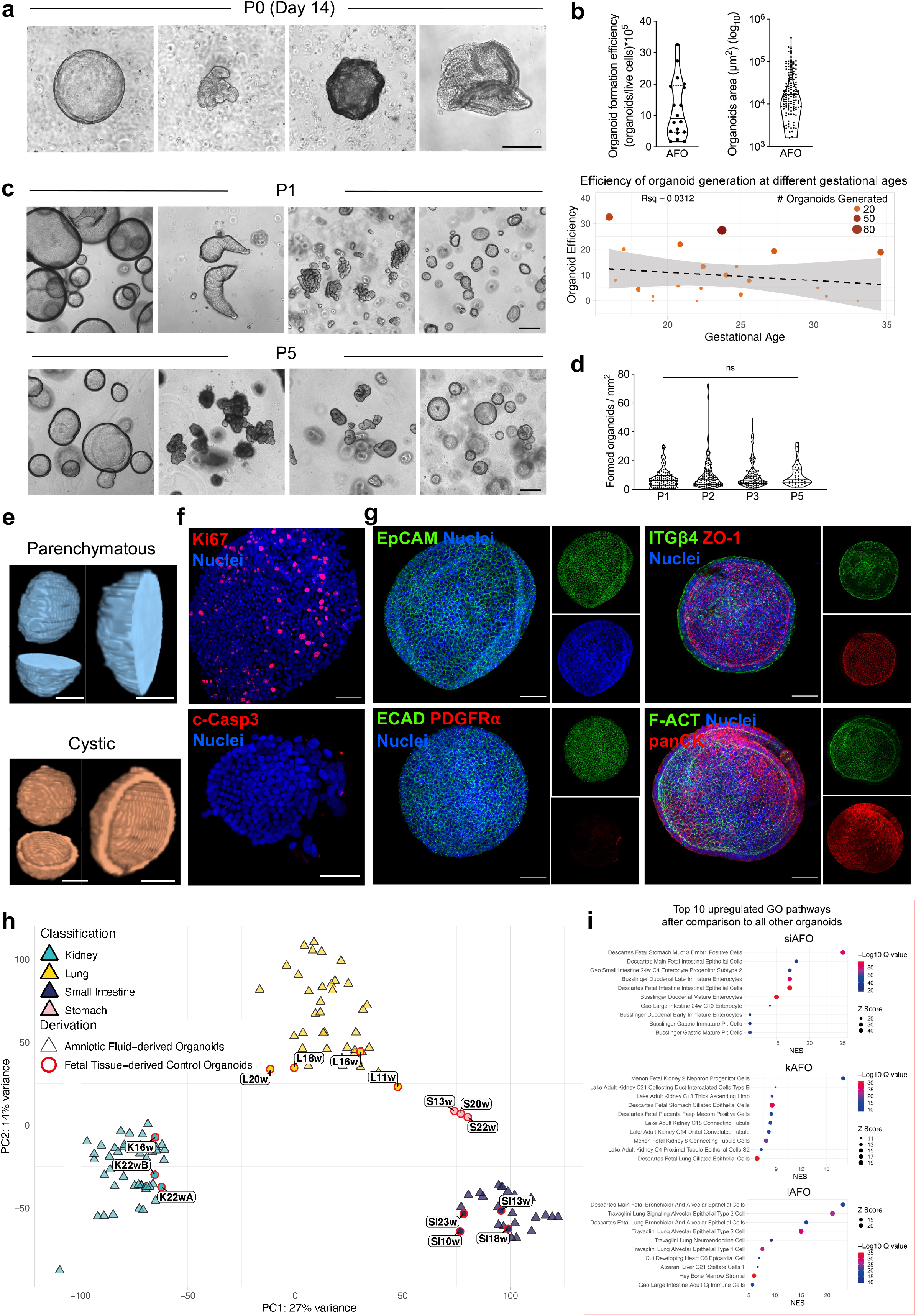
– Generation of primary epithelial fetal AFO. (**a**) Phase contrast images showing the formation of the organoids starting from 3D cultured viable AF cells. The growth of full-size organoids of different morphologies can be observed at day 14 (scale bar: 200 μm). **(b)** Analysis of the formation efficiency (formed organoids/viable cells plated) and of the size (area) of the organoids at isolation (Passage 0) (n=18 AF samples analysed for the efficiency plot and n=128 organoids for area plot; median and quartiles). Dot plot also shown representing organoid generation efficiency (organoids formed / live cells plated) at various gestational ages, with colour and size representing total number of organoids generated per sample. R^2^ = 0.03 and standard error shown. **(c)** Phase contrast images showing the multiplicity of morphologies presented by clonal AFO in expansion, from Passage 1 to 5 (scale bars: 200 μm). **(d)** Quantification of the formed organoids per field of view at 7-15 days of culture quantified over 5 passages (ns: non-significant; *n* =33 organoids from ≥8 AF samples, median and quartiles; 2-way ANOVA with multiple comparison). **(e)** X-ray phase contrast computer tomography (PC-CT) of two of the organoids’ phenotypes observed, parenchymatous and cystic (scale bars: 25 μm). **(f)** The immunofluorescent staining shows expression of the proliferative marker Ki67 and lack of apoptotic cells stained with cleaved caspase 3 in the AFO at P3. Counterstain with Hoechst was used to highlight the localisation of the cell nuclei. (Scale bars: 50 μm). **(g)** Whole mount immunofluorescent staining showing AFO at P3 expressing the epithelial markers EpCAM, ECAD and pan-cytokeratin; together with the lack of the mesenchymal marker platelet-derived growth factor receptor ɑ (PDGFRɑ); the immunofluorescent staining also shows AFO’s polarisation, highlighted by the presence of the epithelial tight junction protein zonula-occludens 1 (ZO-1) on the luminal surface, and integrin beta 4 (ITGβ4) on the basolateral layers. Actin filaments are also displayed by staining with Phalloidin (F-ACT). **(h)** Principal component analysis plot showing the unsupervised clustering of the organoids into three main clusters (n=105 organoid lines from n=17 AF samples). These cluster show co-localisation with primary fetal tissue-derived control organoids (n=11) produced from Lung (Yellow), Small Intestine (Cyan) and Kidney (purple) biopsies. **(i)** Gene ontology analysis of top 10 upregulated pathways in each cluster compared to rest of the dataset, after removal of the fetal tissue-derived controls.

### Generation and maturation of small intestinal Amniotic Fluid Organoids (siAFO)

Small intestinal AFO (siAFO) expanded consistently for over 10 passages, showing the formation of crypt-like structures (Supplementary Figure 3a). Moreover, an EdU incorporation assay indicated the presence of proliferating cells at the base of the intestinal organoids’ crypts (Figure 3a). RNA sequencing of 23 siAFO showed expression of typical intestinal stem/progenitor cell genes (*LGR5, OLMF4*, *LRIG1*, *SMOC2*) as well as Paneth (*LYZ*), goblet (*MUC2, CLCA1*) and endocrine (*CHGA*) cell markers. Enterocyte cell markers (*ALPI*, *FABP1*, *VIL1*, *EZR*, *KRT20, ATP1A1*) were also highly expressed by siAFO (Figure 3b). We validated the RNA sequencing data at a protein level via immunostaining for the crypt stem cell marker Olfactomedin 4 (OLFM4) and the intestinal epithelial cytokeratin 20 (KRT20), highlighting the presence of a crypt-villus axis. Interestingly, siAFO express markers of numerous intestinal cell types such as Paneth cells, as shown by immunostaining for Lysozyme (LYZ), and enterocytes stained for Fatty Acid Binding Protein 1 (FABP1) (Figure 3c and d). siAFO also lack lung and kidney specific markers NKX2-1 and PAX8, respectively (Supplementary Figure 3b). To evaluate the functional capacity of these siAFOs, we assessed digestive enzyme activity. We demonstrated dipeptidyl peptidase IV activity (Figure 3e), a small intestinal brush border enzyme, indicating that siAFOs are capable of peptide hydrolysis. We then performed a maturation assay by placing the siAFO in an intestinal specific medium, in which, upon long-term culture (passage 9) and maturation, siAFO displayed more budding structures, acquiring the typical small intestinal crypt-like organisation. We observed several Chromogranin A-positive enteroendocrine cells, as well as the strong presence of the intestine-specific goblet cell marker Mucin 2 (MUC2) compared to siAFO in the expansion medium (Figure 3f; supplementary figure 3c). In addition, RT-qPCR analysis, following gamma-secretase inhibitor (DAPT) treatment, demonstrated downregulation of both notch target genes (*HES1, OLFM4)* and stem cell, Paneth cell and Wnt target genes (*LGR5, LYZ, AXIN2*). Enterocyte and goblet cell markers (*FABP1, ALPI, MUC2*) were upregulated. Surprisingly, with notch inhibition we did not see an increase in the expression of the master transcription regulator *ATOH1* nor its downstream target *DLL1*. (Supplementary figure 3d)

**Figure 3.**
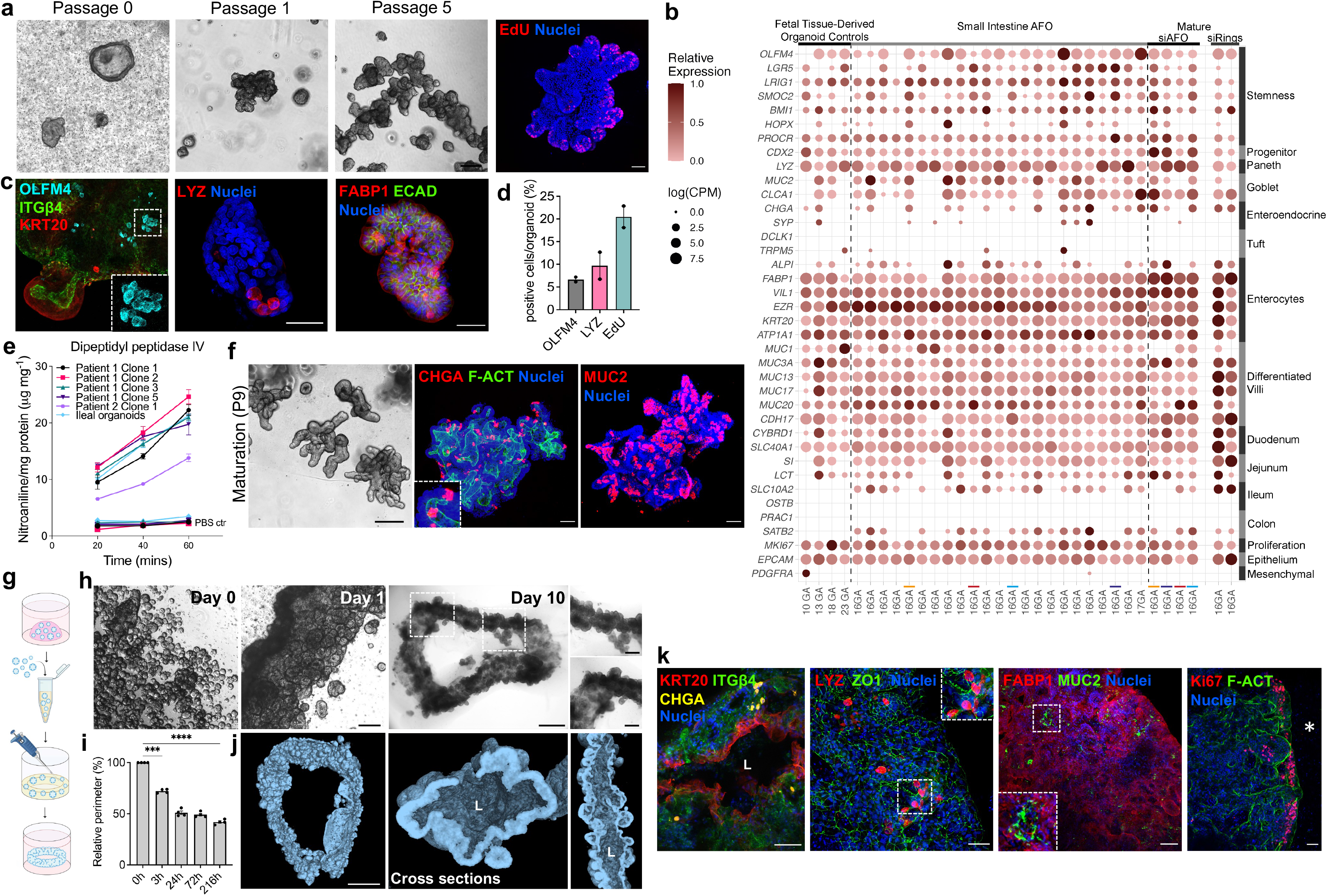
– Generation and maturation of small intestine AFO. (**a**) Phase contrast images depicting the expansion of SIAFO from passage 0 to passage 5 (scale bar: 200 μm). The immunofluorescent staining for EdU (P5) shows the localisation of proliferating cells at the basis of crypt-like structures (scale bar: 50 μm). **(b)** The dot plot shows the gene expression analysis performed on the small intestinal organoids using bulk RNA sequencing showing the presence of all the major small intestine markers in the siAFO (n=2 patients, n=4 siAFO lines in intestinal maturation medium, n=2 siAFO rings, n=4 fetal-derived small intestinal organoids as control). **(c)** Immunofluorescent staining for the intestinal crypt stem cell marker olfactomedin 4 (OLFM4), villi enterocyte marker cytokeratin 20 (KRT20) and luminal integrin-β4 (ITGβ4). The presence of Paneth cells and enterocytes is also highlighted in the immunofluorescent staining for lysozyme (LYZ), epithelial cadherin (ECAD) and fatty acid binding protein 1 (FABP1) respectively; (scale bar: 50 μm). **(d)** Quantification of positive cell type-specific markers siAFO (n=2 independent biological samples; ≥4 organoids per sample; mean ± SEM). **(e)** Functional assessment of Dipeptidyl peptidase IV enzyme in the siAFO; nitroaniline production was monitored within 1 hour of exposure to Gly-Pro p-nitroanilide hydrochloride (n=2 independent biological samples; n=5 clonal organoid lines at passages 6 and 10; n=1 pediatric small intestinal ileal organoid as a positive control; mean ± SEM). **(f)** Phase contrast image depicts the siAFO after maturation, showing budding morphology (scale bar: 200 μm). The immunofluorescent stainings confirm the progression of the maturation by the appearance of Chromogranin A (CHGA)-positive enteroendocrine cells and Mucin 2 (MUC2)-positive secretory cells. Counterstaining with F-actin (F-ACT) and Hoechst shows organoids’ lumen and nuclei respectively (scale bar: 50 μm). **(g)** Schematic depicting the small intestinal ring formation assay. **(h)** The phase contrast images show the formation, self-assembly and compaction of the siAFO ring (scale bar: 200 μm, 1 mm and 500 μm for insets) (n=8 rings from 2 patients). This is also quantified in the bar graph in **(i)** that shows the relative perimeter of the ring decreasing over time (n=4 independent experiments; mean ± SEM; ****P <0.0001 with unpaired t-test). **(j)** 3D reconstructions and cross-sections of a whole siAFO ring following micro-CT scan showing the presence of luminal structures (L; scale bar: 50 μm). **(k)** Immunofluorescent staining on 3D siAFO conducted following tissue clearing confirming the presence of a lumen (L) and KRT20 positive villi-like structures; ITGβ4 highlights the basal side of the intestinal ring together with CGHA positive enteroendocrine cells; the second image displays the presence of lysozyme (LYZ)-positive cells and zonula occludens-1 (ZO-1) in the intestinal ring; mature enterocytes (FABP1) as well as MUC2 secretory cells are present in the SIAFO ring; proliferating Ki67 stained cells are displayed in the crypt-like domain (* external ring side) (scale bar: 50 μm).

Finally, to investigate the potential of siAFO for tissue engineering, we performed an intestinal ring formation assay as previously described^43^ (Figure 3g). After 3 hours from seeding, siAFO started fusing; by day 3 the organoids had self-organised into a more complex tubular and budding structure with full ring maturation at day 10 (Figure 3h-i). RNAseq of siAFO rings revealed a broad, and strong upregulation of genes typical of differentiated and functional intestinal cells, particularly of the mucin secretory lineages, brush border enzymes and enterocytes (Figure 3b). Micro computed tomography (μCT) imaging revealed the self-organisation of the siAFO rings, forming a tube-like structure with a lumen and buds resembling the intestinal architecture (Figure 3j; Supplementary figure 3e; Supplementary Video 1). Importantly, siAFO rings manifested correct cell polarity (KRT20 and ITGβ4) and tight junctions (ZO-1), as well as the presence of enteroendocrine cells (CHGA), Paneth cells (LYZ), enterocytes (FABP1 and KRT20) and goblet secretory cells (MUC2). SiAFO forming intestinal rings also maintained their proliferation capacity on the crypt-like portion of the ring, as shown by the expression of Ki67 (Figure 3k; Supplementary Figure 3f; Supplementary Video 2).

### Generation and differentiation of kidney tubule Amniotic Fluid Organoids (kAFO)

Similar to what has been presented for siAFO, we expanded, characterised and differentiated kidney tubule AF-derived organoids (kAFO), identified by RNAseq within the kidney PCA cluster. Upon expansion, kAFO manifested a more compact morphology distinguishable from the one observed for the siAFO. kAFO could be cryopreserved and expanded long-term (up to passage 10) while maintaining proliferation ability, as highlighted by the diffuse expression of Ki67 (Figure 4a; Supplementary Figure 4a). We then probed our RNAseq dataset for the presence of renal markers, which were present in a total of 44 organoids from 16 patients spanning 18-34 weeks GA. kAFO express canonical renal epithelial development/progenitor and nephron progenitor-specific genes (*PAX2*, *PAX8*, *LHX1, JAG1*). We also detected high expression levels of distal tubule genes (*PCBD1*, *SLC41A3*, *POU3F3*) as well as some proximal tubule markers (*ABCC1*, *ABCC3*, *ABCC4*, *CUBN*). Collecting duct marker *GATA3* was also detected, while canonical Loop of Henle marker *UMOD* was not present. Podocyte markers (*WT1*, *NPHS1, NPHS2*), except for *PODXL,* were not expressed in kAFO. Interestingly, some kAFO lines expressed ureteric bud marker (*RET*), while early cap mesenchyme cell genes (*SIX2*, *CITED1, GDNF*) were not observed (Figure 4b; Supplementary Figure 4b). Based on this profile, we concluded that kAFO manifest a tubuloid-like phenotype and are rich in markers belonging to multiple segments of the renal tubules. As additional validation, we performed immunofluorescent staining to confirm protein expression for the renal epithelium progenitor markers PAX8 and LHX1. Moreover, kAFO displayed kidney segment-specific protein markers such as GATA3 and ECAD (distal tubule/collecting duct), and *Lotus tetragonolobus* lectin (LTL, proximal tubule). The presence of polarised tubular microvilli was also confirmed by immunofluorescence of acetylated tubulin (Ac-αTUB; Figure 4c-d). Interestingly, kAFO exhibited a mixed tubular phenotype with some organoids co-expressing GATA3 and LTL or presenting only GATA3 (Supplementary figure 4c). Functional assessment of kAFO was performed evaluating thallium intake. Following addition of a voltage-gated potassium (K^+^) ion channel stimulator, kAFO showed increased intracellular thallium fluorescence compared to positive control fetal kidney organoids (FKO) and negative control fetal lung organoids (FLO), indicating presence of functional potassium channels (Figure 4e). To further confirm the renal phenotype of the organoids, we have adapted a previously reported differentiation assay to promote maturation of the distal/collecting duct lineage^44^. After 14 days of stimulation with vasopressin and arginine aldosterone, kAFO manifested a slight morphological change and expressed markers of the principal cells of the collecting duct (AQP2) and of the distal tubules (SLC12A1 and CALB1) compared to kAFO in expansion medium (Figure 4f; supplementary figure 4d). Moreover, differentiated kAFO displayed a higher percentage of CALB1 positive cells (19.3 ± 7.6) which was reflected in an increased *CALB1* expression (Figure 4g-h).

**Figure 4.**
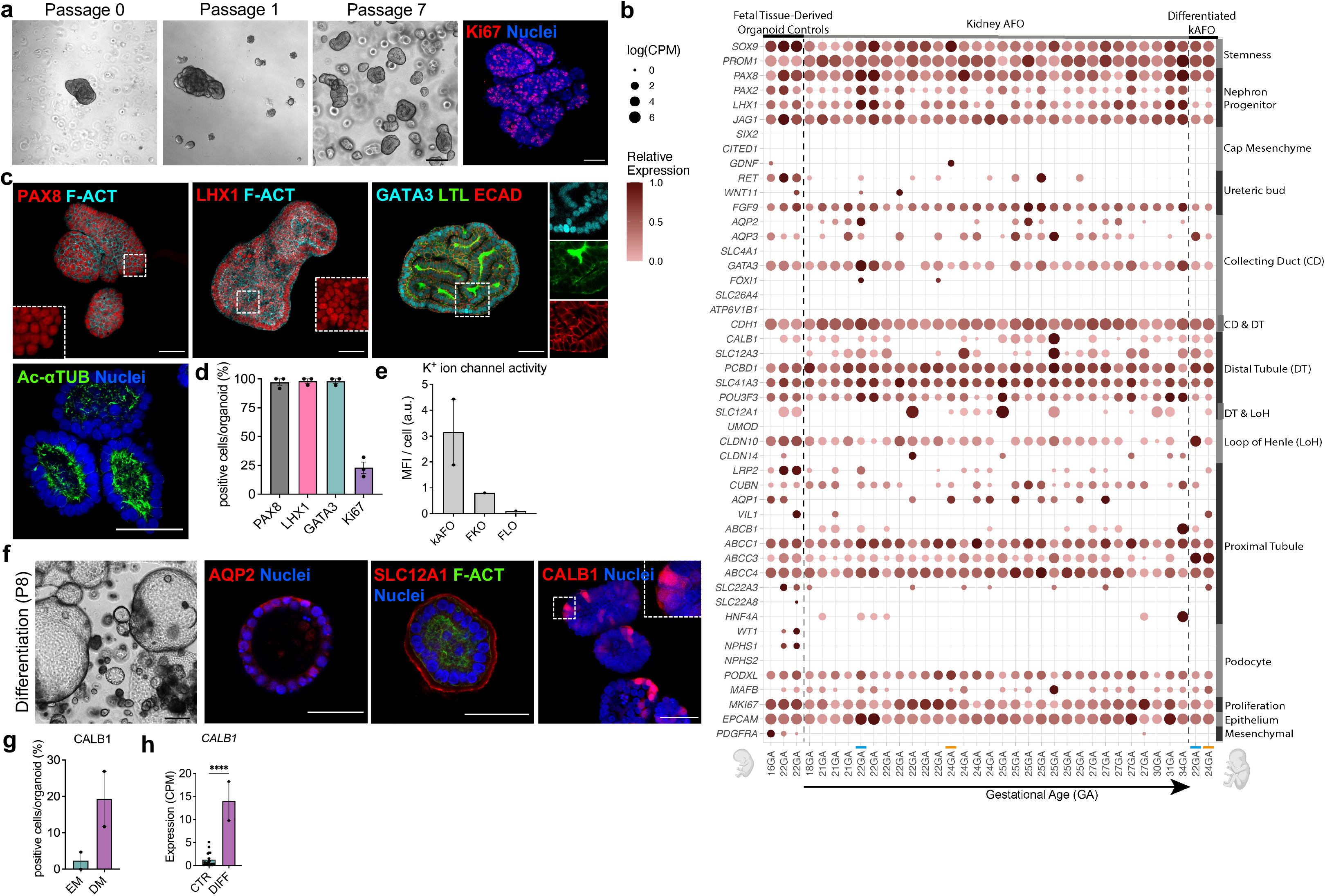
– Generation and differentiation of kidney AFO. (**a**) Phase contrast images depicting the establishment of KAFO starting from passage 0 up to long-term expansion at passage 7 (scale bar: 200 μm). The immunofluorescent staining on the right highlights the presence of the proliferative marker Ki67 (scale bar: 50 μm). **(b)** The dot plot shows the RNAs sequencing analysis showing the presence of a broad range of kidney markers in the kAFO (n=16 patients, n=2 lines in differentiation medium, n=3 independent fetal kidney-derived organoids as a control). **(c)** Immunofluorescent staining shows presence of developmental kidney nephron progenitor markers PAX8 and LHX1, counterstained with F-actin (F-ACT). The panel also shows positivity for the distal tubule/collecting duct marker GATA3, for the proximal tubule marker LTL, as well as the presence of ECAD and apical Ac-αTUB-positive (acetylated α-tubulin) cilia confirmed the renal epithelial identity of the kAFO. (Scale bar: 50 μm). **(d)** The bar graph shows quantification for the early developmental renal markers PAX8, LHX1, and GATA3 in kAFO (n=3 independent biological samples, ≥4 organoids per sample; mean ± SEM). **(e)** Potassium ion channel assay was performed on n=2 kAFO independent biological samples, n=1 fetal kidney organoids, n=1 fetal lung organoids as negative control; medium fluorescence intensity was calculated. **(f)** Phase contrast image and immunofluorescent staining highlighting the morphological changes, as well as the expression of the mature renal markers AQP2, SLC12A1 and CALB1 in kAFO exposed to a renal differentiation medium (scale bars: 200 μm and 50 μm). **(g)** Quantification of CALB1 positive cells in kAFO cultured in expansion medium (EM) compared to differentiation medium (DM) (n=2 biological samples, mean ± SEM). **(h)** Bar graph showing *CALB1* gene expression of differentiated kAFO compared to undifferentiated controls (CTR) based on the RNAseq plot presented in b (n=2 differentiated independent biological samples; mean ± SEM; ****P<0.0001 with unpaired t-test).

### Generation and differentiation of lung Amniotic Fluid Organoids (lAFO)

The lungs are one of the major cellular contributors to the AF due to the continual release of TF, rich in pulmonary cells into the amniotic cavity. Consequently, we hypothesised that lung AF-derived organoids (lAFO) would be readily formed. lAFOs can have a cystic appearance with a thin epithelial cell layer and a lumen or can present with a more parenchymatous morphology. We isolated and clonally expanded 69 lAFO lines from 18 AF samples spanning 16-34 weeks GA, that propagated for over 6 passages maintaining high proliferation ability, as confirmed by the strong expression of Ki67 (Figure 5a; Supplementary Figure 5a-b). RNAseq-based marker analysis indicated the presence of multiple cellular identities within the 38 sequenced lung organoids, with consistent expression of stem/progenitor cell markers (*NKX2-1*, *FOXA2*, *SOX2*, *SOX9*, *TP63, GATA6*), as well as of both Alveolar Type 1 (*HOPX*, *PDPN*, *AGER, AQP5*) and Alveolar Type 2 cells-related genes (*SFTPA1*, *SFTPA2*, *SFTPB*, *SFTPC*, *SFTPD*, *ABCA3*, *LAMP3*). Mature basal cell markers *TROP2* and *NGFR* were not detected when lAFO were cultured in expansion medium. The distal marker *KRT5* was occasionally detected in some lines, as well as the specific ciliated cell transcription factor *FOXJ1,* which showed sporadic low expression. In addition, early secretory cell marker *SCGB3A2* was highly expressed in comparison to the mature marker *SCGB1A1,* which was not expressed. *MUC5AC* secretory cells were present in the control fetal lung organoids, while lAFO did not express the marker. Lastly, neuroendocrine cell marker *ASCL1* was present but not highly expressed by lAFO while few lines showed low expression of *CHGA* (Figure 5b; Supplementary Figure 5c). We then performed protein expression validation via immunostaining, which showed homogenous and strong presence of the stem cell markers NKX2-1, SOX2, and the basal cell marker P63. The prosurfactant protein C (SFTPC) was instead absent (Figure 5c, quantification in 5d). The potential of the lAFO to undergo terminal proximal and distal lung differentiation was also assessed. Remarkably, when pushed towards proximal differentiation, we observed the appearance of a polarised epithelium with motile cilia on the luminal surface of the organoids (Supplementary Video 3). Immunofluorescent staining on the proximally differentiated lAFO showed the presence of Ac-αTUB positive cilia on the luminal side of the organoids, confirming both differentiation and polarisation of the epithelia. The nuclear expression of the ciliated epithelia marker FOXJ1 further corroborated our observation (Figure 5e). In addition, the appearance of the mature basal cell marker keratin 5 (KRT5), and secretory marker mucin (MUC5AC), concomitantly with the maintenance of SOX2, indicated the active proximal lung differentiation process (Figure 5e; Supplementary Figure 5d). Indeed, the percentage of FOXJ1 positive cells (60.7% ± 13.7) was higher in the differentiated lAFO compared to controls in expansion, while the number of KRT5-positive cells (3.1% ± 2.6) did not significantly increase in number (Figure 5f). Moreover, proximal lAFO led to an increased expression of airway markers such as *FOXJ1, TUBA1A* and *KRT5* (Figure 5g). Respiratory motile cilia were analysed in detail by transmission electron microscopy (TEM; Figure 5h). Cilia displayed normal rootlets and associated mitochondria. The ciliary axonemes also displayed normal internal structures such as radial spokes and a normal central microtubule pair. The axonemes were normal in structure, showing outer and inner dynein arms (Figure 5i; Supplementary Figure5e). On the other hand, when pushed towards a distal phenotype, lAFO showed increased protein expression of the AT2 cell marker SFTPB, presenting with different cellular localisation in independent lAFO lines. We observed different distribution of SFTPB in different lAFO lines. Some presenting granules, while others had accumulation of surfactant droplets in the lumen, possibly indicating a higher state of maturation (Figure 5j; Supplementary Figure 5f). This observation was however not reflected in an associated increase in *SFTPB* gene expression (Figure 5k). However, ultrastructural analysis revealed that distalised lAFO contain lamellar bodies with a normal structure and a core composed of multi-lamellar membranes, typical features of distal lung cells (Figure 5l).

**Figure 5.**
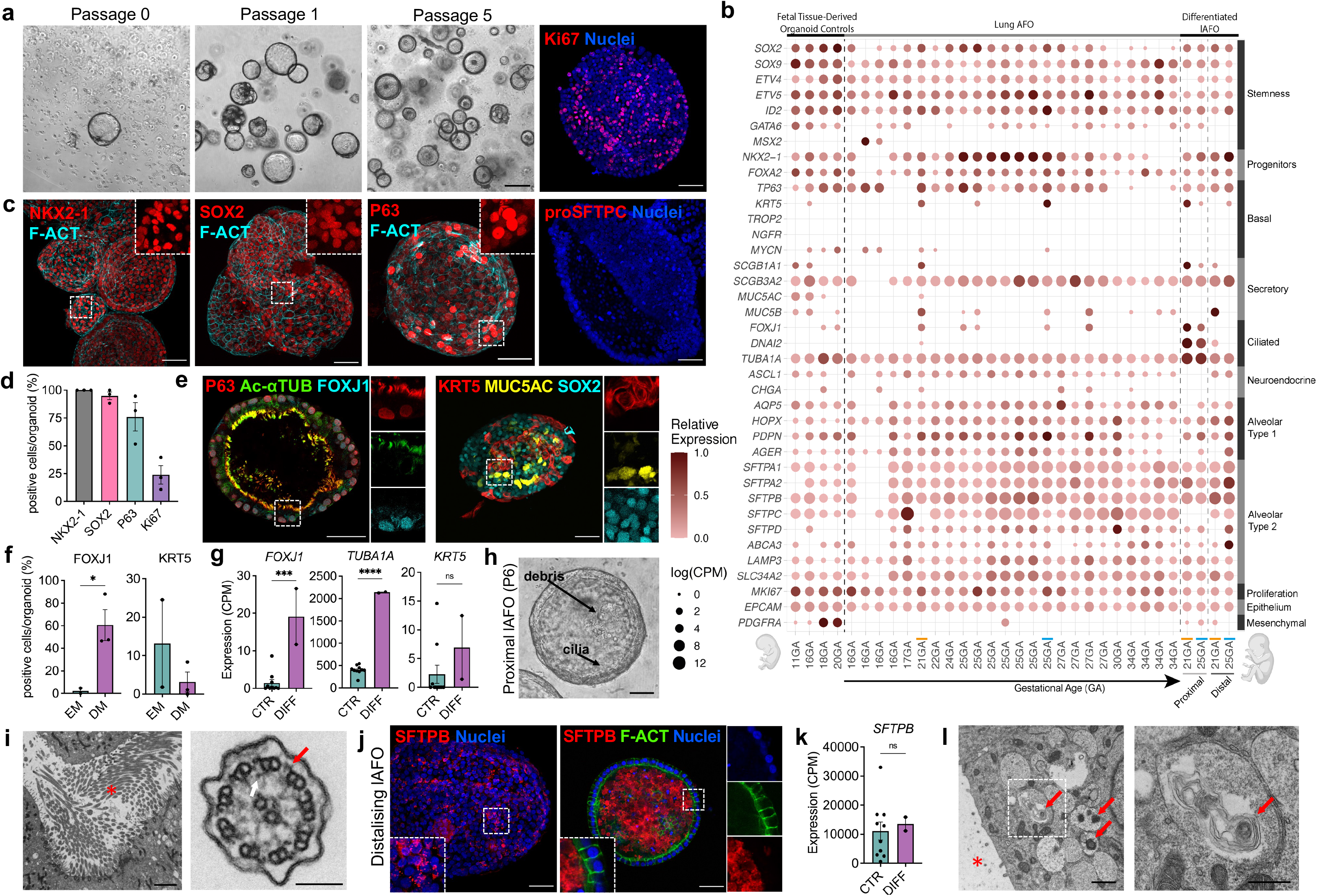
– Generation and differentiation of lung AFO. (**a**) Phase contrast images depicting the establishment of lAFO from passage 0 up to long-term expansion at passage 5 (scale bar: 200 μm). The immunofluorescence on the right highlights the presence of the proliferative marker Ki67 (scale bar: 50 μm). **(b)** The dot plot shows the gene expression analysis performed via RNA sequencing on the organoids (n=18 patients, n=4 differentiated lines, n=4 control tissue-derived primary fetal lung organoids). **(c)** The immunofluorescent stain highlighting the expression of the lung stem/progenitor cell markers NKX2-1 and SOX2. P63-positive basal cells are also present in the lAFO, while proSFTPC secreting cells are absent; images were counterstained with the structural marker F-Actin (F-ACT) and Hoechst, quantified in **(d)** (n=3 biological samples, ≥4 organoids per sample; mean ± SEM) (Scale bar: 50 μm). **(e)** Immunofluorescence staining was performed on lAFO after 14 days of exposure to proximal differentiation protocol. The occurrence of ciliation in the proximally differentiated lAFO is demonstrated by the polarised expression of the ciliary protein Ac-αTUB, co-localised with the basal cell marker P63 and the ciliated cell marker FOXJ1. The panel also shows the expression of the mature basal cell marker KRT5, together with the occurrence of Mucin 5AC goblet cells and maintenance of SOX2 progenitor cells (Scale bar: 50 μm). **(f)** Quantification of FOXJ1 and KRT5 positive cells within the lAFO in expansion (EM) versus proximal differentiation media (DM) (n=3 biological samples, ≥4 organoids per sample; mean ± SEM; *P=0.0461 with unpaired t-test). **(g)** Bar graphs showing gene expression of *FOXJ1*, *TUBA1A* and *KRT5* in proximally differentiated lAFO (DIFF) compared to undifferentiated controls (CTR), based on the RNAseq plot presented in **b** (n=2 biological sample; mean ± SEM; ***P=0.0003, ****P<0.0001 with unpaired t-test). **(h)** Phase-contrast image of proximally differentiated lAFO containing luminal cilia and cellular debris/mucous secretions (scale bar 200 μm). **(i)** TEM imaging showing a cross section of proximal lAFO with cilia inside the lumen (red asterisk) (scale bar 2 μm); In cross-section, the axonemes were normal in structure showing outer (blue arrow) and inner (white arrow) dynein arms (scale bars 100 nm). **(j)** Immunofluorescent staining displaying distalised lAFO showing the occurrence of surfactant-secreting cells (SFTPB); The two images represent two different SFTPB patterning observed in different organoids (granular – left, and luminally secreted – right). Nuclei were counterstained with Hoechst (Scale bar: 50 μm). **(k)** Bar graph showing *SFTPB* gene expression of distalised lAFO (DIFF) compared to control in expansion (CTR) based on the RNAseq plot presented in b (n=2 biological samples; mean ± SEM; ns= nonsignificant). **(l)** TEM of distalised lAFO showing lumen (red asterisk) and cells containing lamellar bodies (red arrows); on the right, magnification of lamellar body containing multi-lammellar membranes (scale bars: 1 μm and 500 nm respectively).

### Lung organoids derived from amniotic and tracheal fluid (AF / TF) of fetuses with CDH manifest a substantially different phenotype compared to gestational age-matched controls

CDH is a congenital malformation where the diaphragmatic muscle fails to close (OMIM: 142340, 222400, 306950), with a consequent herniation of the fetal abdominal organs into the chest. Consequently, the fetal lungs are subjected to a mechanical compression, limiting their physiological growth and leading to developmental impairments of the respiratory and vascular compartments^45, 46^. To investigate the suitability of our platform for fetal disease modelling, we derived lung organoids from both AF and TF of fetuses diagnosed with severe or moderate CDH undergoing fetal intervention at our institutions (Figure 6a; Supplementary Figure 6a; Supplementary Table 1). AF and TF were obtained at the time of Fetoscopic Endoluminal Tracheal Occlusion (FETO)^3, 4^. The fluid collected was subjected to the organoid derivation method described above. As the volume of TF samples collected was low (1-3mL) but with high levels of cell viability (60-75%) we opted to omit the sorting/purification part of the protocol to preserve cell numbers. Similar to the non-CDH lAFO presented in Figure 5, we successfully generated CDH lAFO from 9 CDH fetuses (9/14; 64.3%) and lTFO from 5 (5/9; 55.6%) CDH fetuses (Supplementary Figure 2a). Both CDH lAFO and lTFO expanded for multiple passages (up to P5; figure 6b-d) and expressed NKX2-1, SOX2 and P63 along with Ki67 (Figure 6e). Interestingly, only lTFO expressed SOX9, suggesting a more prominent stem/progenitor identity^25^ (Figure 6f). Moreover, PCA analysis confirmed that all TF-derived organoids had lung identity and demonstrated clustering of CDH lAFO and lTFO with an associated shift of the latter from non-CDH lAFO (Supplementary Figure 6c). We then investigated the gene expression profile of CDH organoids (lAFO and lTFO) using RNAseq and found evidence of expression of most lung epithelial stem/progenitor markers at levels similar to control lAFO (GA-matched; Figure 6g). DEG analysis was performed to explore differences between CDH and control GA-matched lAFO. There was a clear reduction in number of DEGs (p-value < 0.01, |LFC| >2) identified between organoids generated from samples before and after the FETO procedure (181 vs. 47 total DEGs, Figure 6h). Gene ontology analysis identified a clear upregulation in pathways related to surfactant production and metabolism in CDH organoids before FETO treatment vs. age-matched controls, including upregulation of genes related to phosphatidylcholine metabolism (Figure 6i), as well as a downregulation of pathways related to wound healing, growth and differentiation (p-value < 0.05, |LFC| >1). Interestingly, DEG analysis of post-FETO CDH organoids compared to GA-matched controls revealed fewer differences beside surfactant metabolism (Figure 6i; Supplementary Figure 6d). We then subjected CDH lAFO to the same proximal and distal differentiation protocols previously mentioned. Similar to controls, proximal CDH lAFO showed the formation of motile cilia and expressed of acetylated α-tubulin (Ac-αTUB), FOXJ1 and SOX2 proteins (Figure 6j). By recording the ciliary movement using a high-speed camera, we analysed the ciliary beating frequency (CBF) of lAFO from two non-CDH patients and one with CDH. Although remaining within in a physiological range, the CDH lAFO showed a lower CBF of 10 ± 0.17 Hz vs 13 ± 0.16 Hz (Figure 6k; Supplementary Video 4). Finally, when subjected to distal differentiation, CDH lAFO expressed surfactant protein B (Figure 6l), similar to that observed in non-CDH (Figure 5j). We generated lAFO and lTFO from CDH pregnancies from samples taken at the time of the two interventions related to FETO: I) insertion of the occlusive endotracheal balloon (n=9, 26-31 weeks’ GA, n=41, 6 patients); II) removal of the balloon (n=5, 32-34 weeks’ GA, n=13, 3 patients). Paired analysis was not possible owing to sample availability and therefore pooled analysis was performed herein. We noted that there were fewer DEGs in the post-FETO CDH organoids compared to control lAFO of the same gestational age (Figure 6 h, i). Immunofluorescent staining for proSFTPC showed no expression in the CDH lTFO obtained after FETO but was positive in all other CDH samples (figure 6f), highlighting some differences between these samples. Individual comparative DEG analyses between controls, lAFO and lTFO before and after FETO are provided in Supplementary Figure 7.

**Figure 6.**
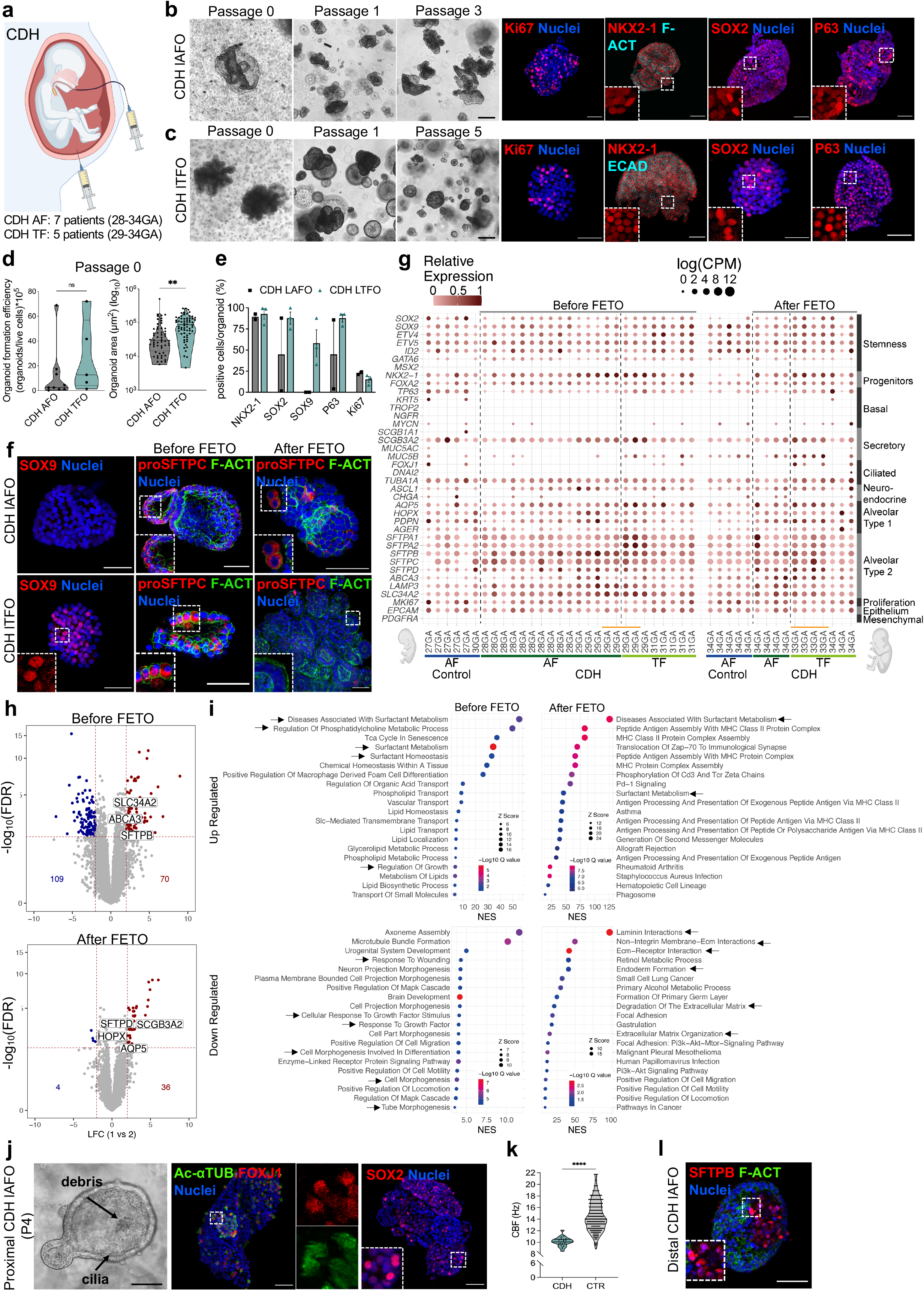
– Generation, differentiation and characterisation of lung AFO and TFO from Congenital diaphragmatic hernia pregnancies. (**a**) Schematic of the amniotic and tracheal fluid sampling from CDH pregnancies. **(b** and **c)** Phase contrast images depicting the establishment of CDH AFO and TFO starting from passage 0 and expanding up to passage 3 and 5 respectively (scale bar: 200 μm). The immunofluorescent staining panel highlights the positivity of the CDH organoids for the proliferative marker Ki67 and the lung stem/progenitor cell markers NKX2-1, SOX2, and the basal cell marker P63 (scale bar: 50 μm); nuclei were counterstained with Hoechst. (**d**) Quantification of the organoids formation efficiency and area of CDH AFO vs. CDH TFO at isolation (n=9 CDH AF and n=5 CDH TF independent samples analysed for the efficiency plot and n≥56 organoids for area plot; median and quartiles; **P=0.0014 with unpaired t-test). **(e)** Quantifications of immunofluorescent stainings are presented in the bar plot (n=2 CDH AF samples, n=3 CDH TF samples, ≥4 organoids per sample; mean ± SEM). **(f)** Immunofluorescent stain showing the expression of SOX9 exclusively in the CDH lTFO. It is also possible to observe the presence of the pro-surfactant protein C (proSFTPC) in both the CDH conditions, and its absence in the CDH lTFO derived after FETO. **(g)** Dot plot showing the expression of lung-associated markers in organoids derived from CDH patients’ tracheal (TF, 29-34 GA, 3 patients) and amniotic (AF, 28-34 GA, 4 patients) fluids alongside GA-matched control lung organoids derived from non-CDH AF (27-34 GA, 3 patients). **(h)** Volcano plots showing differentially expressed genes (DEGs) amongst CDH organoids and GA-matched controls before and after FETO, significant (p-value < 0.01) lung-associated markers are also labelled. **(i)** Pathway analysis showing the 20 top up– and down regulated GO classes in CDH organoids vs. GA-matched controls **(j)** The panel depicts proximal lung differentiation of the CDH LAFO. The phase-contrast image shows proximally differentiated CDH LAFO that display presence of cilia, as highlighted by the immunofluorescence for Ac-αTUB together with the ciliary transcription factor FOXJ1. Nuclei were counterstained with Hoechst (Scale bar for phase contrast: 200 μm; immunofluorescence: 50 μm). **(k)** Quantification of the ciliary beating frequency (Hz) performed using a high-speed camera (n≥22 cilia per organoids, n=1 CDH patient; n=2 non-CDH control patient; median and quartiles, ****P<0.0001 with unpaired t-test). **(l)** The immunofluorescent staining displays the distally differentiated CDH lAFO showing the occurrence of surfactant secreting cells (SFTPB); nuclei were counterstained with Hoechst (Scale bar: 50 μm).

## DISCUSSION

In this article, we describe a reliable method for derivation of autologous primary fetal organoids from multiple epithelial tissues using AF and TF sampled for clinical diagnostic and therapeutic purposes, while allowing the continuation of pregnancy. To guide our experiments, we generated an scRNAseq atlas of the human AF from 11 patients (15 to 34 GA), a resource not currently available in the literature. While it is known that most cells present in the AF are epithelial in nature, their cellular origin has been long debated, and often ascribed to the skin, kidneys and fetal membranes^47^. Our AF cell map confirms that human AFECs are heterogeneous in nature, originating from multiple tissues and consequently distinct from the previously reported placental-derived hAECs^48, 49^. In detail, we provide evidence of the presence, within the AF epithelial cluster, of amniotic fluid-derived stem / progenitor cells of intestinal, renal and pulmonary origin. Once cultured in permissive conditions, AFEC demonstrated their capacity to generate primary clonal epithelial organoids. AFO are amenable to long term expansion, through methodologies similar to that described for fetal organoids produced with destructive approaches. When placed in conditions favourable to promote maturation, the renal, intestinal and pulmonary AFO successfully acquired differentiation hallmarks typical of their tissue of origin.

The generation of AFO uses widely available samples, requires minimal manipulation, and applies only routine organoid culture techniques. The timeline from fluid sampling to full characterisation and expansion of the organoids is currently below 4 weeks, providing a tool that can be applied in a timeline relevant to prenatal counselling and potential therapy, compared with iPSC-dependent methods^16^. The advancement of non-invasive prenatal diagnostic techniques, such as the detection of cell free DNA of fetal origin in the maternal blood stream, has reduced the need to perform amniocenteses for primary diagnostic purposes. However, AF samples remain accessible, with approximately 30,000 amniocenteses performed each year in the UK^50^. This remains necessary in order to perform advanced diagnostics such as array analysis or whole exome sequencing and as confirmatory diagnosis ^2^. Moreover, invasive procedures such amniodrainage, are routinely used as treatment for polyhydramnios^51, 52^, and laser treatment for Twin-to-Twin Transfusion Syndrome (TTTS)^53^. Finally, spina bifida repair, and FETO for CDH, provide further access to the fluid during pregnancy^3–5^.

Intestinal organoids have been isolated from both fetal and adult tissues and used for modelling intestinal development, regeneration, and repair^54–56^. Interestingly, once pushed towards maturation, siAFO acquired features observed in small intestinal organoids derived from primary tissues^56^. Derivation of small intestinal organoids was rare, with success in only two of our samples (16 and 17 GA), one of which was obtained from a termination of pregnancy. The occurrence of AFEC with intestinal stem cell features is an interesting finding given the widely held presumption that after the breakdown of the anal membrane, the anal sphincters retain fetal intestinal content within the gastrointestinal tract during normal development from 12 weeks GA^57^. However, colonic mucosal cells are present in second trimester AF^58^ indicating that cells originating from the gastrointestinal tract are present in AF after the time at which the anal sphincters are thought to retain fetal intestinal content. Thus, there may be the possibility of generating siAFO for therapeutic use in instances where prenatal diagnosis of a congenital disorder potentially resulting in short-bowel syndrome has been made.

In contrast, kAFO and lAFO were easily derived in large numbers across all gestational ages studied. This is presumably related to the regular circulation of fluid through the fetal lungs and renal tract. AF volume increases until the 34^th^ week of pregnancy and then remains relatively constant until term. The production rate of the fetal urine in the human fetus at term (800–1200ml/day), is sufficient to completely replenish the entire amniotic volume every 12– 24 hours ^59, 60^. In keeping with the latter, mesenchymal cells of renal origin have been found in third-trimester AF but are generally difficult to isolate^61^. So far, there has been only one report of isolation of potential podocytes^62^. When cells isolated from kidney tissue and urine are maintained in culture, they are able to generate tubuloids^63^. Tubuloid AFO recapitulate some of the characteristics observed previously, such as the expression of both developmental kidney nephron progenitor markers and, upon differentiation, specific functional proteins.

Similarly, while fetal breathing movements are associated with inward and outward movements of fluid through the trachea, there is overall a net efflux of lung fluid which it is likely to continuously provide lung progenitors to AF, while limiting the influx of cells from other origins into the fetal^64^. This flow is interrupted during FETO treatment and could explain some of the differences between lAFO and lTFO^59^. CDH represents a condition that is clinically relevant for study on account of reliable prenatal diagnosis, along with current clinical practice beginning to offer prenatal therapy for high-risk infant^3, 4^. Current stratification of patients relies on prenatal imaging parameters which are simple to acquire and reliable; although development of sophisticated MRI techniques such as 3-dimensional Lung Volume hold some promise to improve prognostic^65^. The FETO procedure is proven to be of considerable survival benefit to the most severely affected patients; however, there is the need to identify predictive functional biomarkers, as survival remains poor. Notably, around 60% of severe patients do not survive to discharge from hospital^66^.

Previous work on tracheal fluid samples taken before and after FETO showed a different miRNA signature in patients who responded to the intervention, with higher expression of miR-200^67^. This specific pathway could not be robustly analysed in our dataset, since our organoid media contained both A83-01 and Noggin, respectively inhibiting TGFβ and BMP signalling, previously suggested to be an underlying basis for the success of FETO. Importantly, AFO/TFO-derived from fetuses affected by CDH manifest some features of this condition such as alterations in the expression of the surfactant protein genes (SFTPA1, SFTPA2, SFTPB, SFTPC, SFTPD) as presented in Figure 6i. Moreover, increased AT2/AT1 has been suggested as a signature for hypoplastic and CDH lungs in previously published preclinical work^68, 69^. Indeed, our RNAseq analysis demonstrated higher expression of AT2 genes in a greater proportion of cells in the CDH organoids compared to their age-matched controls, with marked upregulation of gene pathways of surfactant metabolism evident on pre– FETO lAFO vs. controls. This suggests that lAFO may be used for disease modelling, drug testing and potentially therapeutically. If successfully implemented, these complex disease models would be amenable for severity prediction, enabling informed clinical decisions to be made before the baby is born. Ultimately, when derived from pregnancies affected by a congenital condition (CDH), AFO and TFO manifested an overall altered phenotype, that was, in part, brought closer to the control transcriptome, in samples taken after treatment with FETO, the current gold standard prenatal approach for this condition (Figure 6h; Supplementary Figure 7).It must be noted that, AF sampling post-FETO is conducted before the removal of the balloon, so AFO may not reflect the changes undergone by the fetal lung in response to the FETO procedure, while TFO being sampled from the occluded trachea will do so^70^. While sampling of TF at the time of FETO is a less widely applicable means of accessing fetal cells the CDH TF samples giving rise to organoids (5 out of 9), only originated lung organoids (5 out of 5) confirming the single tissue origin of the cells contained in the TF.

With rapid advances in organoid biology, particularly with regards of organoids derived from fetal tissue, we expect that in the near future, it will be possible to generate organoids from other tissues exposed to AF. Moreover, we expect that further work will allow different types of organoids to be obtained for each organ. As an example, it is possible to observe the broad distribution of the lung organoids in the PCA presented in Figure 2, as well as the formation of two renal clusters. The core advantage of the technology presented in this article, is to offer the ability of deriving fetal organoids prenatally, without the need for accessing the fetal tissue. Currently, the only viable alternative is derivation of organoids from iPSC. Although the iPSC route offers flexibility, and virtually unlimited expansion potential, it comes with a series of technological trade-offs prevent widespread clinical use. Some of these issues are intrinsic to reprogrammed cells, such as insertional mutagenesis of the reprogramming transgenes, acquisition of genomic aberrations during expansion, aneuploidy, sub-chromosomal copy number variants point mutations and alteration of epigenetic marks^71^. Moreover, the use of mutated iPSCs, could influence their successful recapitulation of disease phenotypes^72^. From our perspective one of the major technological burdens of using reprogramming to produce fetal organoids is the time required. If the aim is to model or treat a condition before birth in order to provide personalised prognostic information or autologous organoid-based therapy, it must be possible to implement this strategy within the 40 weeks of gestation. This timeline is further shortened by the fact that AF / chorionic villi sampling is not normally performed until the end of the first trimester; and furthermore, the majority of prenatal therapies are currently ideally delivered prior to 30 weeks. Several examples of iPSCs-organoid derivation highlight the need of at least 21 weeks to produce the organoids^73^. Our approach benefits from the already committed progenitors present in the fetal fluids, which require minimal manipulation to lead to the production of a large amount of primary autologous fetal organoids in approximately 4 to 6 weeks.

### Limitations

One consideration of the work presented here relates to the availability of samples for research. The invasiveness of amniocentesis and its associated risks to the pregnancy means it is performed less frequently, and extremely rarely on pregnancies deemed low risk for congenital anomalies. In the context of the current study, samples were taken in the course of clinical work-up which makes the contextualisation of the phenotype observed in tissue-specific AFO challenging. Certainly, this is an issue that would need to be addressed in disease-specific manner in the future. For example, we would not expect fetuses with identified congenital heart disease to necessarily have issues affecting their pulmonary epithelia and as such these could provide relevant control data for CDH.

Moreover, the sorting strategy used in this work was based on the use of Hoechst and PI. We recognise that these dyes (particularly Hoechst) are not ideal for the subsequent culture of cells in a translational setting. Follow-up studies will investigate the use of alternative non-toxic / non-intercalating dyes.

The initial characterisation of the organoids presented in the current study has been conducted using RNA sequencing. The high running cost of this technique can be problematic, in particular when trying to apply this technology on a large scale. However, we believe that this could be replaced with more cost-effective methods such as PCR validation, immunohistochemical staining or image-based characterisation. Notably, the use of TF organoids would not be affected by this problem as all TFO formed with a lung identity. Amongst over 200 organoids lines that were sequenced, we found 3 to have a hybrid renal/pulmonary phenotype (Supplementary figure 2d). This has been ascribed to a non-clonality of the initial organoid at picking stage. Despite the efforts in plating the cells at low density for the primary culture, this may have been the results of a doublet, or the fusion of two nascent organoids. This is however a low-occurrence phenomenon (3 lines out of the 159 studied) and can be addressed by future studies implementing appropriate QC.

More generally, our system is limited to the epithelial compartment. Despite aligning with most of the reports on primary organoids, our system will not allow in its current form the modelling of complex conditions involving, for example, the mesenchymal and vascular compartments. In our opinion, this could be overcome by culturing the organoids in combination with mesenchymal, endothelial cells. Importantly, mesenchymal cells have broadly been reported in the AF^30, 33^. Moreover, these cells have been successfully directly reprogrammed to the endothelial lineage^74, 75^, suggesting the amenability of different AF cell lineages to complex prenatal disease modelling.

## Conclusions

In conclusion, we report derivation of epithelial organoids of different tissue identity through a minimally invasive approach, from continuing pregnancies and within a broad GA window. AFO of intestine, kidney and lung origin are expandable and can be functionally differentiated with great potential for functional diagnosis, regenerative medicine, and disease modelling. An example of the latter could be observed in CDH, where AFO can recapitulate some of the original features observed in the affected fetuses, suggesting their use for personalised medicine.

## Acknowledgements

We are thankful to Alessandro Pellegata, Marco Pellegrini, Valerija Karaluka, Robert Hynds, Daniyal Jafree, David Long, Sara Mantero, Nicola Elvassore, Carla Pou Casellas, Anna Baulies, Hans Clevers and Muzlifah Haniffa for the useful discussions and advice. We are grateful to Ayad Eddaoudi and the GOSICH Flow Cytometry facility, Dale Moulding and the GOSICH Imaging facility, Tony Brooks and UCLGenomics. We thank the University of Leicester Core Biotechnology Services Electron Microscopy Facility for help with material processing. Control fetal tissues for this study were provided by the Human Developmental Biology Resource. We are thankful to the UCL Hospital (UCLH) FMU research midwife team, Roland Devlieger, Luc De Catte, Liesbeth Lewi, Sofia Mastrodima-Polychroniou, and Mr George Attilakos who helped collecting the fluid samples. We thank the Diamond Light Source for the provision of beam time. Finally, we are grateful to the Human Developmental Cell Atlas network, whose members provided valuable advice across the development the project.

## MATERIALS AND METHODS

### Amniotic fluid collection and isolation of the viable cell fraction

AF samples (amniocenteses and amniodrainages) were collected from the University College London Hospital (UCLH) Fetal Maternal Unit (FMU) and UZ Leuven as part of standard patient’s clinical care (REC 14/LO/0863 IRAS 133888). Written informed consent was obtained prior to the procedure. After collection, fluids were stored at 4°C until processing. AF samples were passed through a 70 μm and 40 μm cell strainer and transferred in 50ml tubes before being centrifuged at 300 g for 10 min at 4°C. Supernatant was discarded, pellet resuspended in 5-10 mL of FACS blocking buffer containing 1% FBS and 0.5 mM EDTA in PBS and transferred to FACS tubes. Cells were incubated with 5 μg/mL Hoechst (Sigma-Aldrich, 33342) for 40 min at 37°C and then counterstained with 2 μg/mL propidium iodide (PI) (Sigma-Aldrich, P4170) for 5 min at RT. Viable cells were sorted using a FACSAria III (BD), unselected for side and forward scatter, but gated for Hoechst^+^ and PI^-^.

### Derivation and culture of human amniotic fluid organoids (AFO)

Viable amniotic fluid cells were resuspended in 30 μL of Matrigel (Corning) and plated 6×10^5^ live cells/droplet onto a pre-warmed 24 well plate. Cells were cultured in an ad hoc defined generic medium (Supplementary Table 2) supplemented with Rho-kinase inhibitor (ROCKi; Tocris) and Primocin. To establish clonal organoid lines, single organoids formed were manually picked at passage 0 under the microscope to be clonally expanded. Each organoid was transferred in a 0.5 mL tube pre-coated with 1% BSA (Sigma-Aldrich). Organoids were resuspended in TrypLE (Thermo) and incubated for 5 min at 37°C. After digestion, organoids were disaggregated by pipetting and additional 400 μL of ice-cold Advanced DMEM/F-12 supplemented with Glutamax, P/S and Hepes (ADMEM+++) were added. Organoids were precipitated with a minicentrifuge for 2 min and a second washing passage was repeated. After centrifugation the pellet was resuspended in 20 μL Matrigel (Corning) and plated in a 48 well plate. The plate was incubated for 20 min at 37°C and generic medium was added with ROCKi (Tocris) and Primocin for the first 3 days. Medium was replaced every 3-4 days. After approximately 10 days, grown organoids were passaged as described below.

### Derivation and culture of human tracheal fluid organoids (TFO)

Tracheal fluids (TF) were collected during procedures of Fetoscopic Endoluminal Tracheal Occlusion (FETO) carried out at the University College London Hospital (UCLH) or KU Leuven, kept refrigerated and processed within 24-48 hours. We collected TF samples before the insertion of the balloon and after its removal. Due to the nature of the TF samples, mostly small and containing a majority of living cells, FACS sorting was not performed. TF was transferred into a 15 mL tube on ice, washed with ice-cold ADMEM+++ and centrifuged at 300 g for 5 min at 4°C. Supernatant was discarded and cells were resuspended in 1 mL of ADMEM+++. Cells were counted and plated in Matrigel droplets. Plates were incubated for 20 min at 37°C and human fetal lung organoid medium (Supplementary Table 2) supplemented with ROCKi and Primocin was added. Medium was changed every 3 days. TFO were clonally expanded and passaged as described below.

### Passaging of organoids

Depending on number and size, organoids were passaged to a 24 or 12 well plate after clonal expansion. Afterwards organoids were usually split 1:2 to 1:3 after 10–14 days of culture. The medium was aspirated and ice-cold ADMEM+++ was added to each well. Matrigel droplets were disrupted and collected into a 15 ml tube on ice. Organoids were washed with 10 mL of cold ADMEM+++ and centrifuged at 300 g for 5 min at 4°C. Big and cystic organoids were resuspended in 1 mL of ADMEM+++ and mechanically disaggregated using a P1000 pipette. If small, organoids were disrupted enzymatically. Medium was aspirated and organoids pellet was resuspended in 300 μL of TryplE. After incubation for 5 min at 37 °C, organoids were pipetted with a P200 to break them down into single cells. Cold ADMEM+++ was added up to 10 mL and the sample was centrifuged at 300 g for 5 min at 4°C. Supernatant was discarded and cell pellet was resuspended in Matrigel and plated. The plate was incubated for 20 min at 37°C to allow the Matrigel to solidify, upon which generic culture medium was added with ROCKi. Medium was changed every 3 days.

### Fetal tissue samples collection and derivation of control primary fetal organoids

Control fetal tissue samples were sourced via the Joint MRC/Wellcome Trust Human Developmental Biology Resource under informed ethical consent with Research Tissue Bank ethical approval (Project 200478: UCL REC 18/LO/0822 – IRAS ID 244325; Newcastle 18/NE/0290 – IRAS ID 250012). The derivation of control fetal organoids was conducted as follows:

#### i) Human fetal small intestinal organoids

Fetal small intestines were processed as previously described^76^. Tissue was washed with PBS, cleared of any mesenteric tissue and fat, then cut longitudinally. The villi were scratched away using a glass coverslip. The remaining tissue was cut into 2–3 mm pieces, washed vigorously, and incubated in 2 mM EDTA in PBS for 30 min for 5 min on an orbital shaker. The supernatant containing the intestinal crypts was centrifuged at 800 g for 5 min at 4 °C. After being washed in ADMEM+++ and centrifuged, the pellet was resuspended in Matrigel and plated in presence of Primocin and ROCKi. Medium recipe is in Supplementary Table 2.

#### ii) Human fetal kidney tubular organoids

The process was adapted following a previously published protocol for deriving adult tubuloids^77^. Briefly, fetal kidneys were harvested, washed in ice-cold HBSS and minced to isolate the cortical tissue. This was washed in 10 mL of basal medium and supernatant was removed when the tissue pieces were sedimented. After being washed several times in ADMEM+++, tubular fragments were isolated by 1 mg/mL collagenase digestion (C9407, Sigma) on an orbital shaker for 30-45 min at 37°C. Fragments were further washed in basal medium with 2% FBS and centrifuged at 300 g for 5 min at 4°C. Pellet was resuspended in Matrigel and cultured in Kidney medium (Supplementary Table 2) supplemented with ROCKi and Primocin.

#### iii) Human fetal lung organoids

Fetal lung tissue was processed adapting a previously published protocol^20^. Briefly, fetal lungs were minced and washed in ADMEM+++. Tissue fragments were digested in ADMEM+++ containing 1 mg/mL of collagenase (C9407, Sigma) on an orbital shaker at 37°C for 30-60 min. The digested tissue was shacked vigorously and strained over a 100 μm filter. Tissue fragments were washed in ice-cold basal medium with 2% FBS and centrifuged at 300 g for 5 min at 4°C. Supernatant was discarded and pellet was resuspended in Matrigel and cultured in Lung medium (Supplementary Table 2) supplemented with ROCKi and Primocin.

#### iv) Human fetal stomach organoids

Fetal stomach organoids were isolated from specimens following an established dissociation protocol^24^. Briefly, stomachs were cut open and mucus was removed with a glass coverslip and mucosa was stripped from muscle layer. Mucosa samples were cut into pieces of 3-5 mm and washed in HBSS until the supernatant was clear. Tissue was incubated in chelating buffer supplemented with 2 mM EDTA for 30 min at RT. Tissue fragments were squeezed with a glass slide to isolate the gastric glands which were transferred in ADMEM+++, strained through at 40 μm and centrifuged at 300 g for 5 min at 4°C. Pellet was resuspended in Matrigel and plated. Gastric medium (Supplementary Table 2) was added with ROCKi and Primocin.

### Organoid cryopreservation and thawing

After 7-10 days of culture, organoids were dissociated enzymatically as described above. The final cell pellet was resuspended in 1:1 ADMEM+++ and freezing medium (80% FBS and 20% DMSO). Cryovials were stored at –80°C overnight and then transferred to LN2 for long term storage. For organoids thawing, cryovials were equilibrated on dry ice and then placed at 37°C. Vial content was rapidly transferred to 15 mL falcon tube containing 9 mL of ice-cold ADMEM+++, then centrifuged at 300 g for 5 min at 4°C. Supernatant was discarded and pellet resuspended in cold Matrigel (Corning). After 20 min of incubation at 37°C, medium was supplemented with ROCKi and replaced after 3 days.

### Evaluation of organoids formation efficiency and area

Organoid formation efficiency was determined by counting the number of organoids at passage 0. The total number of formed organoids per well was manually counted approximately 14 days after seeding of the AF cells in Matrigel. The efficiency was determined by calculating the total number of grown organoids divided by the number of viable single cells initially plated. The organoids area was determined by measuring the perimeter of each organoid in different 5x fields acquired at the Zeiss Axio Observer A1 and using ImageJ software and normalised by field size.

### Organoid maturation / differentiation

Small intestinal AF organoids (siAFO): after manual passaging, organoids were seeded in triplicate in Matrigel and cultured in generic medium. After approximately 7 days, human small intestine medium was used (Supplementary Table 2) for 14 days. Basal culture medium was the same but without addition of CHIR99021, DAPT (notch inhibitor) 10μm was added to basal culture medium for 48 hours to stimulate differentiation.

Kidney AF organoids (kAFO): after either manual or enzymatic passaging, organoids were seeded in triplicate in Matrigel and cultured in generic medium. After approximately 7-10 days, distal/collecting duct kidney differentiation medium (Supplementary Table 2) was used for 14 days.

Lung AF organoids (lAFO): after manual passaging, organoids were seeded in triplicate in Matrigel and cultured in generic medium for approximately 10 days. For lung proximal differentiation, PneumaCult™ ALI Medium (StemCell Technologies, #05001) was then used for 14 days. For distalisation, previously reported medium^78^ was used for 14 days as well (Supplementary Table 2).

### Intestinal ring formation in Collagen hydrogel

siAFO were expanded for 7 days in generic medium. Matrigel droplets were dissolved in Cell Recovery Solution (Corning, 354253) for 45 min at 4°C. The organoids were collected and resuspended in 120 μL of Collagen hydrogel (Collagen type I 0.75 mg/mL, DMEM-F12 1X, HEPES 1M, MilliQ to volume, pH 7) and plated in an ultra-low adherent 24 well plate making a circular shape around edges of the well. Plate was incubated for 30 min at 37°C and rings were cultured in intestinal medium for 10 days.

### Whole-mount immunofluorescence

Prior to fixation, organoids were removed from Matrigel using Cell Recovery Solution for 45 min on ice. Organoids were harvested into a 15 mL tube pre-coated with 1% BSA in PBS and fixed with 4% PFA for 20 min at RT. Samples were washed 3 times with PBS for 5 min and spun down at 300 g for 5 min at 4°C. Whole-mount immunostaining was performed by blocking and permeabilising the organoids with PBS-Triton X-100 0.5% with 1% BSA for 1 hour at RT. Primary antibodies were incubated in blocking/permeabilisation buffer for 24 h at 4 °C in rotation. After being extensively washed with PBS-Triton 0.2%, organoids were incubated with secondary antibodies overnight at 4 °C in rotation. Organoids were further washed and resuspended in PBS for confocal imaging. For tissue clearing of siAFO ring, previously published protocol was adapted^79^. EdU staining was performed with the Click-iT EdU Alexa Fluor 568 Imaging kit (Life Technologies) following the manufacturer’s protocol and images were acquired using a Leica SP5 confocal microscope. Full list of antibodies is available in Supplementary Table 3.

### Image acquisition

Phase-contrast images were acquired using a Zeiss Axio Observer A1. Immunofluorescence images of whole-mount staining and sections were acquired on a Zeiss LSM 710 confocal microscope using 25x, 40x, and 63x immersion objectives. Image analysis and z-stacks projections were generated using ImageJ (Rasband, W.S., ImageJ, U. S. National Institutes of Health, Bethesda, Maryland, USA, https://imagej.nih.gov/ij/, 1997-2018).

### X-Ray phase contrast computed tomography

The imaging of the organoids was performed using phase contrast computed tomography (PC-CT) at beamline I13-1 (coherence branch) of the Diamond Light Source (Didcot, UK). The x-ray energy was 9.7 keV and the system resolution 1.6*μm*. The organoids were imaged embedded in Histogel (Epredia™ HistoGel™). The PC-CT scan entailed the acquisition of 2000 equally spaced projections through a 180° rotation of the specimen. The total scan time was approximately 1h. The “single image” phase retrieval operation^80^ was applied to the acquired projections, with the estimated phase to attenuation ratio (refer to as δ/β ratio) set at 250. Both phase retrieval and tomographic slice reconstructions were performed using Savu^81^ whilst the 3D images were generated using Drishit^82, 83^.

### Micro-focus computed tomography imaging

After 10 days of culture, AF intestinal rings were collected and processed for micro-focus computed tomography (micro-CT) scanning. Rings were fixed for 1 h in 4% PFA and extensively washed in PBS. The specimen was iodinated overnight by immersion in 1.25% potassium tri-iodide in 10% formalin solution. After being rinsed in deionised water, the sample was wrapped in laboratory wrapping film and mounted in Histogel (Epredia, HG-4000-012) in 1.5 mL tube. Micro-CT scanning was performed using a Nikon Med-X micro-CT scanner (Nikon Metrology, Tring, UK). The specimen was mounted and held in pace using a drill chuck to ensure centralised rotational positioning. Whole specimen scans were acquired using an x-ray energy of 120 kV, current of 50 μA, exposure time of 1000 ms, 4 frames per projection, a detector gain of 24 dB and an optimised number of projections of 2258. A tungsten target was used and an isotropic voxel size of 3.57 μm was achieved. Reconstructions were carried out using modified Feldkamp filtered back projection algorithms with CTPro3D (Nikon, Metrology) and post-processed using VGStudio MAX (Volume Graphics GmbH).

### siAFO dipeptidyl peptidase IV assay

Organoids were plated in 48-well plates, 15μl BME/well, in triplicate. Organoids were washed in PBS and then incubated at 37°C with 200μl /well in Gly-Pro p-nitroanilide hydrochloride (Sigma G0513) dissolved in PBS at a concentration of 1.5mM (or PBS alone in control wells). During incubation, samples were agitated on an orbital shaker (60rpm) and supernatants were sampled at 20, 40 and 60mins. Absorbance (415nm) was measured with a plate reader (Biorad) and concentration determined by comparison to a 4-nitroaniline (Sigma 185310) standard curve (0-200µg/ml) and normalised mg^-1^ organoid lysate protein (Pierce BCA Protein Assay Kit – ThermoScientific).

### kAFO Potassium ion channel assay

kAFOs were expanded at least in triplicate in a 96 well plate. FluxOR™ II Green Potassium Ion Channel Assay was performed according to the manufacturers’ instructions (F20017, Thermo). Briefly, medium was removed and 80 µL of 1X Loading Buffer were added to each well and incubated for 30 min at RT and 30 min at 37 °C to facilitate dye entry. After removing the Loading Buffer, 80 µL of Assay Buffer were added to each well. The plate was read using a microplate reader every 5 sec for 5 min, with an excitation wavelength of 480 nm and an emission wavelength of 545 nm. After 5 min of recording, voltage-gated channels were stimulated with 20 µL of High Potassium Stimulus Buffer containing 2 mM Thallium Sulphate (Tl_2_SO_4_) and 10 mM Potassium Sulphate (K_2_SO_4_). The plate was read once again every 5 sec for 5 min. The average of replicates was normalised on the control (Assay Buffer) and the number of cells.

### Ciliary beat frequency analysis

Organoids were seeded into 8 well glass bottom slide and differentiated towards the lung proximal lineage as described above. For ciliary beat frequency (CBF) analysis, motile cilia grown inside organoids were observed using an inverted microscope system (Nikon Ti-U) with a digital high-speed video camera (Prime BSI Express, Teledyne photometrics). Videos were recorded at a rate of approximately 87 frames/second using a 60x objective. 8-10 videos of individual organoids for each patient were acquired, and the total region of interest (ROI; 1024 x 1024) of each video was then divided into 16 smaller ROIs (sROIs; 256 x 256) for analysis. Every sROI containing clearly visible cilia was counted for 5 full sweeps and the number of frames for 5 full sweeps was integrated into the calculation of CBF (Hz) followed by [87 / (number of frames for 5 sweeps)] x 5.

### Transmission electron micrograph

Organoids in matrix were fixed in a mix of 2.5% Glutaraldehyde and 4% Paraformaldehyde in 0.1M Sorensen’s buffer, pH7.3, and washed with 0.1M Sorensen’s buffer. They were postfixed in 1% aqueous solution of osmium tetroxide, washed and dehydrated through the increasing series of ethanol solutions, followed by propylene oxide (Merck). Organoids were embedded in TAAB812 resin (TAAB Laboratory Equipment Ltd) and sectioned to approximately 70nm thick using a Leica UC7 Ultramicrotome (Leica Microsystems). Sections were collected onto copper mesh grids and contrasted for 2min with 4% uranyl acetate solution in methanol (VWR), followed by 2min in lead citrate (Reynold’s solution). Samples were viewed on a JEOL JEM-1400 TEM (JEOL UK) with an accelerating voltage of 120kV. Digital images were collected with a Xarosa digital camera using Radius software (both from EMSIS).

### Flow cytometry

Cold-stored AF was centrifuged at 300 g for 10 min at 4°C. Supernatant was discarded, pellet resuspended in FBB and strained over a 70 μm and then at 40 μm filter. Cells were incubated with the following fluorochrome-conjugated antibodies: APC/Fire™ 750 anti-human CD324 (E-Cadherin) (Biolegend 324122), PE anti-human CD326 (EpCAM) (324205).

### RNA isolation and RT-qPCR

Organoids were collected from Matrigel with Cell Recovery Solution for 45 min at 4°C. Cells were then washed in ice-cold PBS to remove Matrigel leftover. Organoids were centrifuged at 300 g for 5 min at 4 °C and supernatant discarded. Pellet was resuspended and lysed with RLT buffer (Qiagen). Total RNA was isolated with RNeasy Micro or Mini Kit (Qiagen) following manufacturer’s instructions. RNA concentration was quantified using a Nanodrop (Thermo). cDNA was prepared using High-Capacity cDNA Reverse Transcription Kit (Applied Biosystems, #4368813). Quantitative real-time PCR detection was performed using PowerUp™ SYBR® Green Master Mix (Applied Biosystems, A25742) and StepOnePlus Real-Time PCR System (Applied Biosystems). Assays for each sample were run in triplicate and were normalised to the housekeeping gene β-actin. Primer sequences are listed in the Supplementary Table 4.

### Bulk RNA Sequencing

RNA extraction was performed as described above and stored at –80°C until processing. NEBNext Low Input RNA library preparation and sequencing was performed by the UCLgenomics facility. Single-end bulk RNA sequencing was performed on an Illumina NextSeq 2000. 100 cycles were run to achieve an average of 5million reads per sample.

### Transcriptome bioinformatic analysis

Quality control was conducted on the FASTQ raw sequences using v0.11.9 FastQC (https://github.com/s-andrews/FastQC). v0.6.6 TrimGalore! (https://github.com/FelixKrueger/TrimGalore) was used to trim low quality reads (quality 20, length 70). v2.7.1a STAR (https://github.com/alexdobin/STAR) was applied to align the FASTQ sequences to the NCBI human reference genome GRCh38.p13 (https://www.ncbi.nlm.nih.gov/assembly/GCF_000001405.39/). v1.6.3 featureCounts (https://doi.org/10.1093/bioinformatics/btt656) quantified the expression of individual genes to generate the raw count matrix, using the GRCh38.104 gene annotation (ensembl.org/Homo_sapiens/Info/Index). Default parameters used for both alignment and quantification. The generated count martrix was further processed with a custom R script. Genes will less than 10 reads across 3 samples were removed. Gene IDs were included using the added BioMart package (ensembl.org/info/data/biomart/biomart_r_package.html). Counts per million (CPM) normalisation was completed with the edgeR package (bioconductor.org/packages/release/bioc/html/edgeR.html).

ComBat_seq batch correction (rdrr.io/bioc/sva/man/ComBat_seq.html) was applied between the 4 batches. ggplot2 was used throughout for graph generation, including principal component analyses (PCA) and dot plots generated from the normalised count matrix. Clustered heatmaps were generated with Pheatmap (cran.r–project.org/web/packages/pheatmap/), to enable hierarchical comparisons.

For the comparison of CDH– and healthy-derived AFO, the pheatmap hierarchical clustering was used to separate the samples for differentially expressed genes (DEGs) analysis using DESeq2 (bioconductor.org/packages/release/bioc/html/DESeq2.html). Volcano plot was generated with ggplot2 to highlight statistically significant DEGs.

### Single-cell RNA Sequencing

Library generation was done following the 10x Genomics Chromium Next GEM Single Cell 3ʹ Reagent Kits v3.1 (Dual Index) kit. Libraries were sequenced using NovaSeq 6000. Data processing was completing with CellRanger. Analysis was done with Seurat V4.1.1 within a custom R script was used for major downstream processing. Cells with less than 150, or more than 8000 features were removed to prevent doublets or cells of low quality. Dying cells with more than 30% mitochondrial genes detected were also removed. Normalisation was then carried out using the NormalizeData function, with a LogNormalize method and a scale factor of 10,000. Batch correction was completed through Seurat’s IntegrateData function, after assessing for integration anchors based on the 2000 most variable features. The object was scaled using ScaleData, RunPCA and FindNeighbors determined for 20 PCs. UMAPs and violin plots were generated using ggplot2 and normalised gene expression always shown, with violin plots showing averaged normalised gene expression within the identified epithelial cluster. v1.6.1 SingleR^84^ was used to label a single cell experiment object using Human Primary Cell Atlas Data (humancellatlas.org), accessed via celldex (github.com/LTLA/celldex).

### Statistics and reproducibility

Statistical analysis was conducted on data from at least three independent experimental or biological replicates wherever possible, as stated in the figure legends. Results are expressed as the mean ± S.E.M or as the median and quartiles (25% and 75% percentiles) or range. Statistical significance was analysed using unpaired Student’s *t*-tests for comparisons between two different experimental groups. Statistical significance was assessed using one-way ANOVA with Dunnett’s or Tukey’s post hoc multiple-comparisons test for analysis among more than two groups. \**P* < 0.05, \*\**P* < 0.01, \*\*\**P* < 0.001, \*\*\*\**P* < 0.0001 were considered significant. Exact *P* values are stated in each relevant figure legend where appropriate. Statistical analysis was carried out using R and GraphPad Prism 9 software.

## DATA AVAILABILITY

Raw and processed data of the bulk RNA sequencing data (AFO/TFO) and scRNAseq (AF) have been uploaded to NCBI GEO (awaiting accession number) and will be made publicly available in due course. For now, access keys have been supplied to the editor.

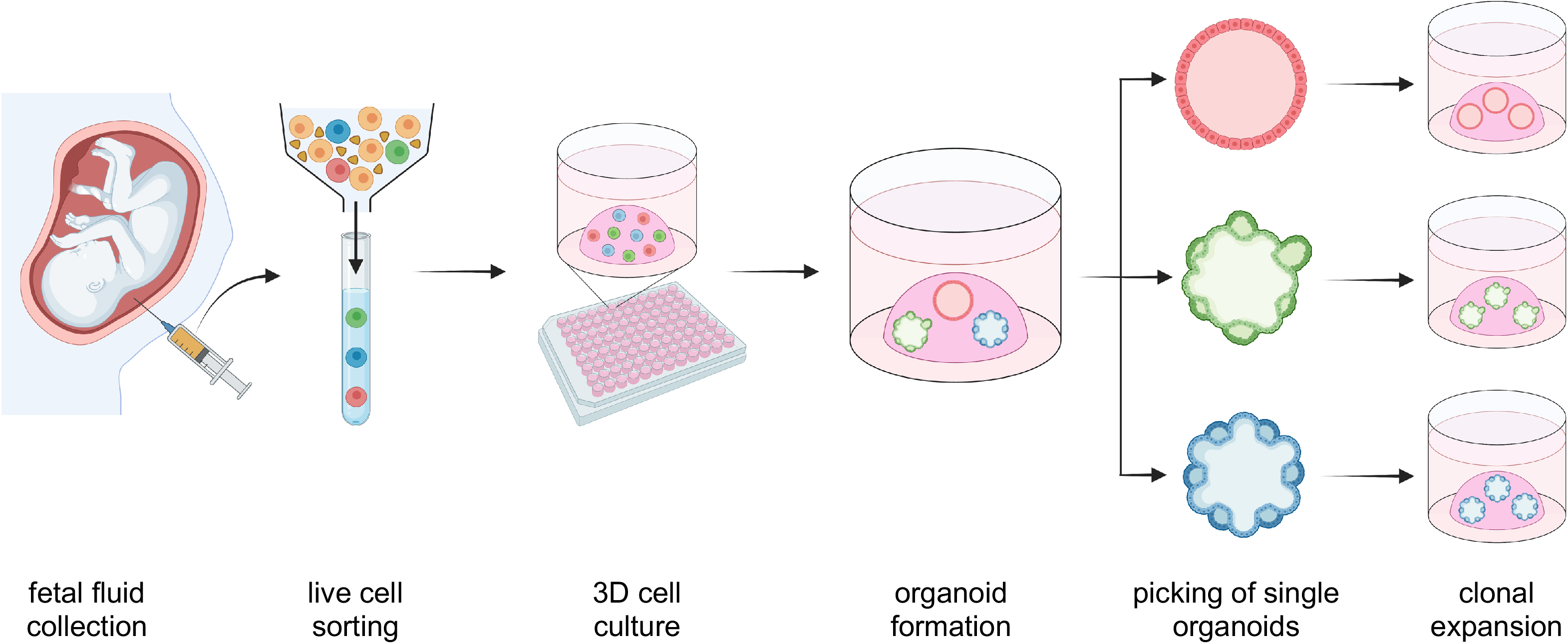

